# A Matter of Degrees: Latitudinal Variation in the Transcriptional Response to High and Low Temperatures in an Estuarine Cnidarian

**DOI:** 10.64898/2026.04.14.718487

**Authors:** Janki A. Bhalodi, Adam M. Reitzel

## Abstract

Populations of the same species inhabiting distinct geographical regions must meet the requirements of local thermal regimes to survive. While individuals integrate both deeply conserved and genotype-specific transcriptional responses to temperature shifts, unique local requirements may diversify the balance between these two mechanisms in distinct populations. The sea anemone *Nematostella vectensis* inhabits highly variable estuarine environments across a broad geographic range, providing an excellent system to investigate how local adaptations shape responses to temperature stress. While studies have explored the genotypic and phenotypic diversity among *N. vectensis* populations, the diversity in transcriptional responses to heat and cold remain poorly understood. We used RNA sequencing to characterize transcriptional programs in *N. vectensis* from Nova Scotia (NS), Maryland (MD), and Florida (FL) under acute temperature treatments at 10°C and 38°C. Individuals exhibited a stronger response at 38°C than at 10°C, with NS and MD responses being similar and FL exhibiting a unique response. A core set of genes was differentially expressed across all populations under heat stress, while responses to cold were highly population specific. To evaluate the role of a key transcription factor, heat shock factor (HSF), we analyzed the presence of HSF binding sites (HSEs) in promoters of differentially expressed genes (DEGs). Upregulated genes containing three or more promoter HSEs were strongly induced at 38°C in MD and FL, but not in NS. To identify the involvement of other transcription factors, we searched for overrepresented motifs in the promoters of the top 100 DEGs at 38°C, revealing a differential enrichment of motifs across the three populations. Together, these findings suggest that *N. vectensis* populations utilize diverse transcriptional programs in response to common hot and cold temperatures.

## 1. Introduction

Global climate change is increasing temperatures in natural habitats, affecting species fitness and ecosystem functioning (Traill, et al. 2010; Pigot, et al. 2023; Wiens and Zelinka 2024). While many species are vulnerable to temperature changes, ectotherm animals, which cannot regulate their internal temperature, are particularly impacted because their biochemical reactions are largely temperature-dependent (Nguyen, et al. 2011; Huey and Kingsolver 2019; Jørgensen, et al. 2022). Species inhabiting highly variable thermal environments also show limited capacity for acclimation and are more susceptible to climate change (Tomanek 2010). A variety of evolutionary conserved molecular and cellular processes (e.g., heat shock response, redox regulation, metabolic and mitochondrial adjustments) are deployed in response to acute and chronic thermal stress. How these processes may vary between sensitive and resistant species or populations of the same species is informative for a better understanding of how differences in genetics may translate to variation in resistance. Therefore, it is critical to study the mechanisms of how ectotherms inhabiting variable environments may acclimate and adapt to changing temperatures.

*Nematostella vectensis*, a sea anemone belonging to the phylum Cnidaria, is a sessile invertebrate native to shallow, brackish estuarine environments. Distinct populations of *N. vectensis* are distributed along the Atlantic and Pacific coasts of the United States, as well as parts of the United Kingdom (Hand and Uhlinger 1994). While populations on the western coast of the US are thought to be introduced and are largely clonal (Darling, et al. 2009), populations in their natural distribution occur along the thermal cline of the eastern coast, subjecting these genetically diverse populations to highly variable temperatures across their range (Reitzel, et al. 2008; Reitzel, Herrera, et al. 2013). These individuals must cope with daily temperature fluctuations of nearly 20 °C and seasonal temperature differences as high as 40 °C within their environments (Reitzel, Chu, et al. 2013). Additionally, even across short geographic distances, populations of *N. vectensis* exhibit high genetic divergence and significant population structure, suggesting limited gene flow and the potential for adaptation to local environments (Reitzel, et al. 2008).

Evidence for local adaptation across geographically distinct populations of *N. vectensis* has been documented in various studies (Friedman, et al. 2018; Sachkova, et al. 2020; Baldassarre, et al. 2023; Smith, et al. 2023). Populations exhibit a negative relationship between growth rate and latitude of origin under elevated temperatures, suggesting that southern populations of *N. vectensis* are locally adapted to their relatively warmer environments (Reitzel, Chu, et al. 2013). Individuals from different locations also differ in oxidative stress resistance that manifest as variations in tentacle numbers, tentacle regeneration, and survival rates following exposure to hydrogen peroxide (Friedman, et al. 2018), which may be related to the distribution of redox-sensitive variants of the transcription factor NF-κB (Sullivan, et al. 2009). Population-specific differences in key life history characters such as venom production in response to heat, salinity, and UV stressors have also been reported in *N. vectensis* (Sachkova, et al. 2020), which may be related to large differences in gene copy number (Smith, et al. 2023; Surm, et al. 2024). Responses to abiotic conditions for *N. vectensis* are also impacted by non-genetic factors, specifically the microbiome. Bacterial community composition varies among different *N. vectensis* genotypes and within the same genotypes at different temperatures, underscoring the role of such adaptations in the stress response (Baldassarre, et al. 2023).

While many studies have identified genotypic and phenotypic diversity across *Nematostella* populations, the diversity of transcriptional responses to thermal stress remain less understood. In *N. vectensis*, thermal tolerance is not only a plastic trait allowing individuals to survive in a broad range of temperatures, but also one with a genetic basis, as offspring from parents from different locations had different temperature tolerances showing differential expression of some general stress response genes (Rivera, et al. 2021). However, the potential genetic differences that account for such differential thermal performance remain unclear. Waller, et al. (2018) showed that a subset of HSP70 genes had generally similar expression under acute and chronic high temperature conditions for multiple populations of *N. vectensis*. Sachkova, et al. (2020) showed differential expression of a subset of HSPs and superoxide dismutase genes in response to increased temperatures (36°C), with some variation among *N. vectensis* populations. These previous studies have largely focused on the transcriptional response to high temperatures for *N. vectensis*. However, the role of the broader transcriptional diversity in contributing to local thermal adaptation, warm and cool, has not been studied in *N. vectensis*.

In this study, we exposed clonal populations of *N. vectensis* from Nova Scotia (NS), Maryland (MD), and Florida (FL) to acute cold (10 °C) or heat (38 °C) stress for three hours. We utilized RNA-seq to quantify the diversity of transcriptional responses to these stressors, allowing us to characterize shared and unique molecular signatures in geographically distinct populations. Our findings highlight a robust transcriptional response to heat stress, identifying a core set of heat-responsive genes with common expression patterns across the three populations. The responses to cold stress were comparatively modest. Each population also had large population-specific transcriptional mechanisms supporting a conclusion that each location deploys unique molecular responses to common temperature changes.

## 2. Materials and Methods

### 2.1 Anemone culturing

Clonal lines of adult *Nematostella vectensis* females from Nova Scotia (Canada), Maryland (USA), and Florida (USA) were maintained in the laboratory at room temperature (∼25 °C) in 15 parts per thousand (PPT) artificial seawater (ASW), prepared using Instant Ocean Sea Salt and ultrapure water. Anemones were given water changes weekly and fed freshly hatched *Artemia nauplii* two to three times per week.

### 2.2 Temperature stress experimental design

To investigate the transcriptional response of each *N. vectensis* population to temperature stress, clonal lines were first acclimated at 25 °C in an incubator without light for two weeks with normal feeding. Three anemones per population were collected at time zero (T0) and transferred to individual 1.5 mL Eppendorf tubes. Immediately, the tubes were submerged in liquid nitrogen to flash-freeze and then stored at -80 °C until RNA extraction. A subset of anemones was moved to glass bowls containing ∼350 mL ASW pre-cooled to 10 °C, pre-heated to 38 °C, or maintained at 25 °C. Each bowl thus represented one condition and contained four replicate anemones each. The bowls were then moved to incubators pre-set to each respective temperature. All treatments exposures were conducted in the dark to minimize the effects of light as a variable. At the 3-hour time point, three anemones per condition were collected and preserved, as described above.

### 2.3 RNA extraction and sequencing

Total RNA was extracted from each replicate anemone per condition using the Qiagen Allprep DNA/RNA kit following the manufacturer’s protocol. Anemone tissue was homogenized for extractions using a syringe until all tissue was dissolved. The Agilent 4200 TapeStation system was used to obtain the concentration and RNA-integrity number (RIN) for each sample. Two samples, one replicate each from the FL T0 and MD T0 conditions, showed low concentrations such that RIN value could not be determined for them; these samples were removed from the study at this stage. All other samples had a RIN value greater than 8 and were used for sequencing. Libraries were generated for each sample using the NEBNext UltraExpress RNA Library Prep kit (NEB #E3330), in conjunction with the NEBNext Poly(A) mRNA Magnetic Isolation Module (NEB #E7490) and NEBNext Multiplex Oligos for Illumina (NEB #E6444) kits, as described by the manufacturer. Paired-end sequencing of pooled libraries was performed by Admera Health using the NovaSeq X Plus platform.

### 2.4 Analysis of differential gene expression and gene ontology terms

Sequence quality-check was performed using FastQC v0.12.1 (Andrews 2010) and MultiQC v1.21 (Ewels, et al. 2016). Reads were trimmed using Fastp v0.22.0 (Chen 2025). Quantification and alignment of reads to the *N. vectensis* transcripts (jaNemVect1.1) (Fletcher and Pereira da Conceicoa 2023) was performed using Salmon v1.10.3 (Patro, et al. 2017). Differential expression analysis was conducted using DESeq2 v1.44.0 (Love, et al. 2014), and the resulting data were normalized using a variance-stabilizing transformation in R. Significantly differentially expressed genes were considered to be those with an absolute log2 fold change value greater than 2 and a padj value of less than 0.05. Venn diagrams were created using the package ggVennDiagram v1.5.4 (Gao, et al. 2021) and heatmaps were generated using the package Pheatmap v1.0.12 (https://CRAN.R-project.org/package=pheatmap). Gene ontology enrichment analysis was performed using the NIH DAVID web server (Sherman, et al. 2022). All R analyses were performed using R v4.4.1 in RStudio. Detailed scripts are available on our GitHub repository (https://github.com/jbhalodi/Temperature-Stress-RNA-Seq).

### 2.5 Promoter motif analyses

The 500bp putative promoter sequences of all *N. vectnesis* genes were extracted from the reference genome assembly (jaNemVect1.1) using a previously published Python script (Cleves, et al. 2020). These sequences were searched for the canonical heat shock element (HSE) sequence ‘GAANNTTCNNGAA’ using the FIMO tool from MEME Suite (https://meme-suite.org/meme/tools/fimo) (Bailey, et al. 2009; Grant, et al. 2011). FIMO output was used to quantify the number of motifs in each gene promoter, and this file was used to visualize the relationship between promoter HSE count and the magnitude of expression of genes in R (see GitHub). Promoters of top differentially expressed genes for each population were additionally searched for up to two enriched motifs using the MEME tool from MEME Suite. Enriched motifs were searched against a database of known transcription factor binding motifs using the TOMTOM tool from MEME Suite.

## 3. Results

### 3.1 Global transcriptomic responses of *N. vectensis* populations to thermal stress

To investigate the transcriptomic changes induced by high and low temperature stress, we performed RNA-sequencing on *Nematostella vectensis* anemones from three populations (Nova Scotia (NS), Maryland (MD), and Florida (FL) following acute temperature treatments at 10 °C (cold stress), 25 °C (control), and 38 °C (heat stress). Principal component analysis (PCA) revealed that the primary source of variation was driven by temperature, with PC1 accounting for 40% of the variance and clearly separating all 38 °C-treated samples from all other temperature treatments (Fig. 1). PC2 represented 29% of the variance, largely separating all FL samples and the NS and MD samples. These data suggest that FL anemones harbored a unique transcriptional signature even at control temperatures, whereas MD and NS anemones exhibited very similar expression profiles across all temperature treatments. Interestingly, we did not observe any considerable variance between the control and 10 °C-treated samples, suggesting that this temperature did not elicit extensive transcriptional changes in anemones from these populations.

**Figure 1.**
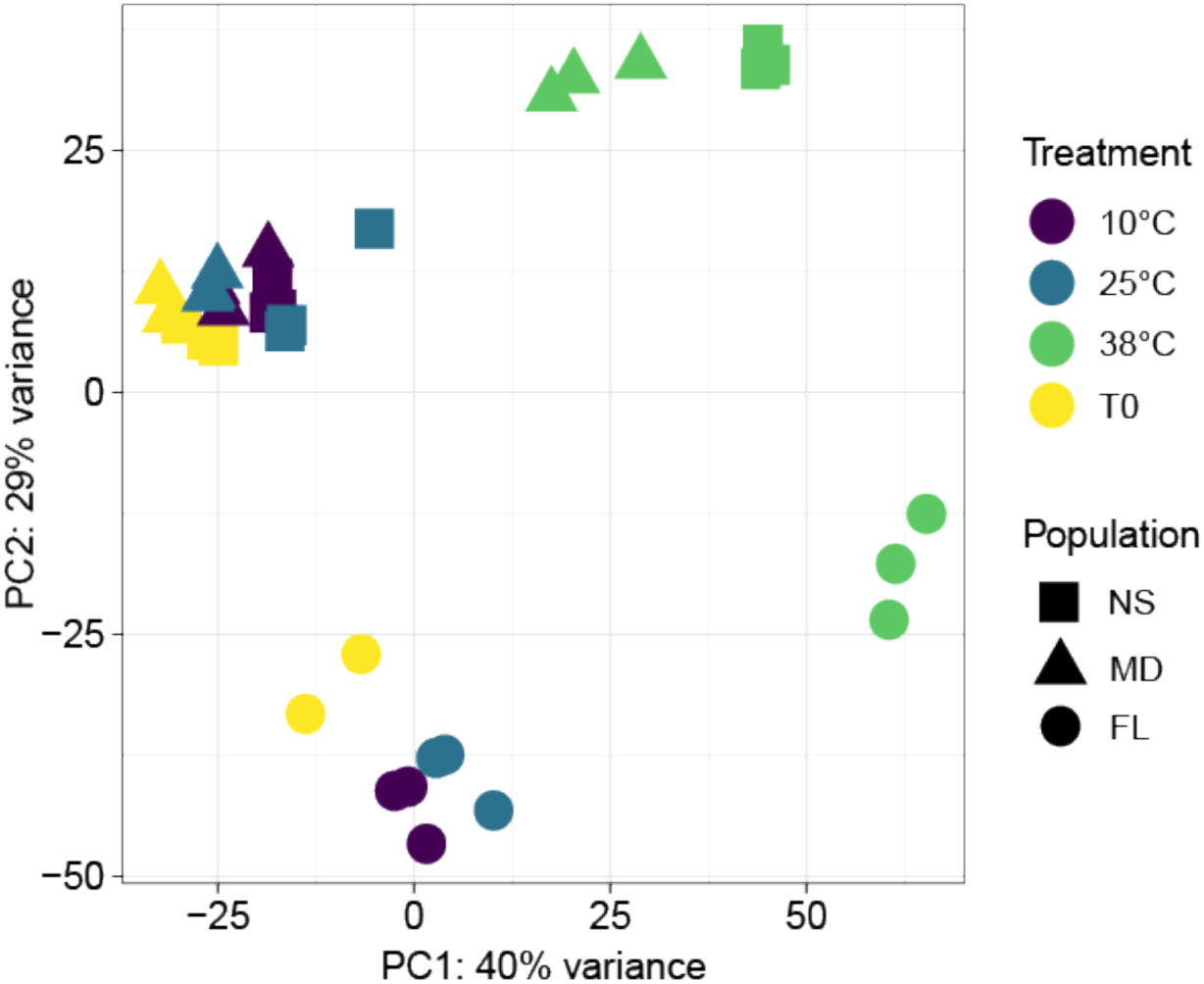
Principal component analysis (PCA) of global expression of *Nematostella vectensis* populations across temperature treatments. Analysis performed on all vst-normalized genes. Percentages represent variance explained by each principal component. Colors represent temperature treatments (°C), where T0 are samples before anemones were transferred to the three temperature treatments. Shapes represent each of the three *N. vectensis* populations studied. NS, Nova Scotia; MD, Maryland; FL, Florida.

### 3.2 Differential expression analysis reveals a large response to heat stress

To identify the molecular drivers of the response to temperature treatments, we performed differential gene expression (DGE) analysis using DESeq2. To ensure that the identified transcriptional changes were specifically driven by temperature responses, we performed a baseline comparison between the 25 °C control and samples collected at timepoint zero (T0). Genes significantly differentially expressed in this comparison were considered artifacts of experimental handling and were excluded from all subsequent analyses of cold and heat stress responses (Fig. S1). Overall, exposure to 38 °C induced a larger transcriptomic shift than exposure to 10 °C across all populations (Fig. 2). Intriguingly, NS exhibited the greatest number of differentially expressed genes (DEGs) under both cold (n = 125) and heat (n = 2539) stress conditions. In contrast, FL showed the fewest DEGs under cold stress (n = 60), while following NS closely in the number of heat stress DEGs (n = 2380). The MD population exhibited a comparable response to cold stress relative to FL (n = 64). However, under heat stress, MD showed the lowest number of DEGs (n = 1613), a count considerably lower than that of the FL anemones.

**Figure 2.**
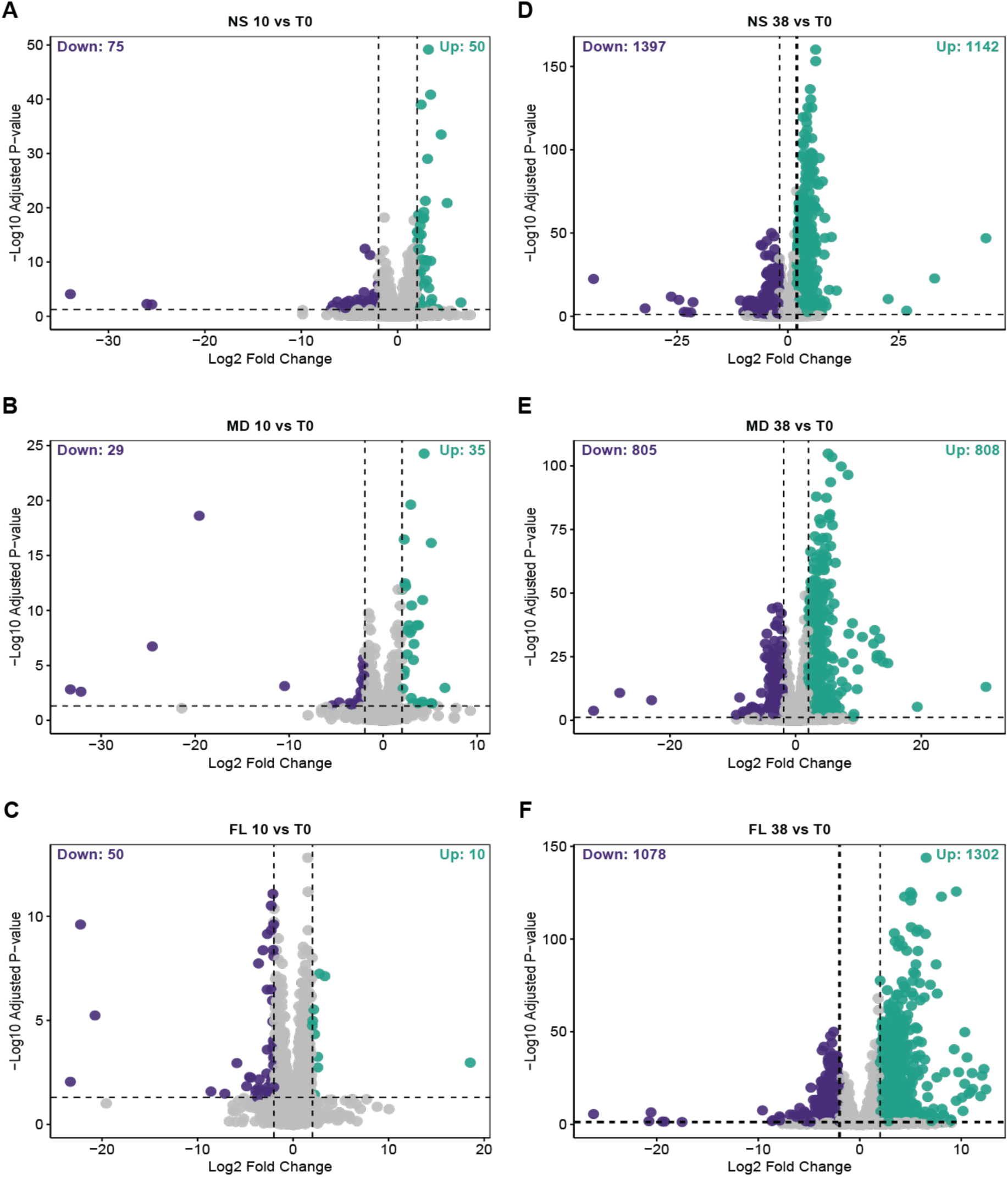
Volcano plots of differentially expressed genes in treatment samples relative to timepoint zero samples (T0). Significantly differentially downregulated genes (log2 fold change < -2, padj < 0.05) in purple (left) and upregulated genes (log2 fold change > 2, padj < 0.05) in teal (right) for each comparison. Cold stress: (**A**) Nova Scotia (NS) 10 °C, (**B**) Maryland (MD) 10 °C, and (**C**) Florida (FL) 10 °C. Heat stress: (**D**) NS 38 °C, (**E**) MD 38 °C, and (**F**) FL 38 °C.

### 3.3 Cold stress elicits a largely population-specific gene expression pattern

Although the magnitude of the cold response was relatively small, functional enrichment analysis revealed that the nature of this response was highly population-specific (Fig. 3A). Highlighting this divergence, only a single significant DEG, the transcription factor *Jun* (5521019), was co-upregulated across all three cold-stress datasets (Fig. 3B). *Jun* is a well-characterized component of the activator protein-1 (AP-1) transcription factor complex, known to play a role in cell proliferation, differentiation, apoptosis, and response to stimulus (Karin, et al. 1997; Shaulian and Karin 2002; Agron, et al. 2017); In *N. vectensis*, this *Jun* has been identified as the cnidarian-specific homolog of c-Jun belonging to the stress-induced AP-1 transcriptional complex (Sunagar, et al. 2018). We also identified only a single shared DEG between the FL and NS populations: a putative skeletal organic matrix protein (125573772), which was downregulated in both populations (Fig. 3C). Homologs of this gene in corals contribute to biomineralization and formation of the coral skeleton, which has protective functions (Drake, et al. 2013). FL shared eight DEGs with MD, which were functionally enriched for protein refolding, response to heat, and cellular response to unfolded protein (Table S1). Interestingly, several of these shared genes exhibited divergent expression patterns between the two populations (Fig. 3D). In contrast, while we identified 15 shared DEGs between NS and MD, no significant functional enrichment was detected for this set; however, these genes showed similar expression profiles in both populations (Fig. S2). Finally, no significant functional enrichment was observed for the DEGs unique to each population. The minimal overlap in DEGs and the lack of a shared functional signature suggest that *N. vectensis* lacks a unified ‘core’ cold-stress program at this threshold. This stands in stark contrast to the response observed at 38 °C, where a robust set of several hundred genes was co-regulated across all populations, representing a conserved emergency response to acute heat.

**Figure 3.**
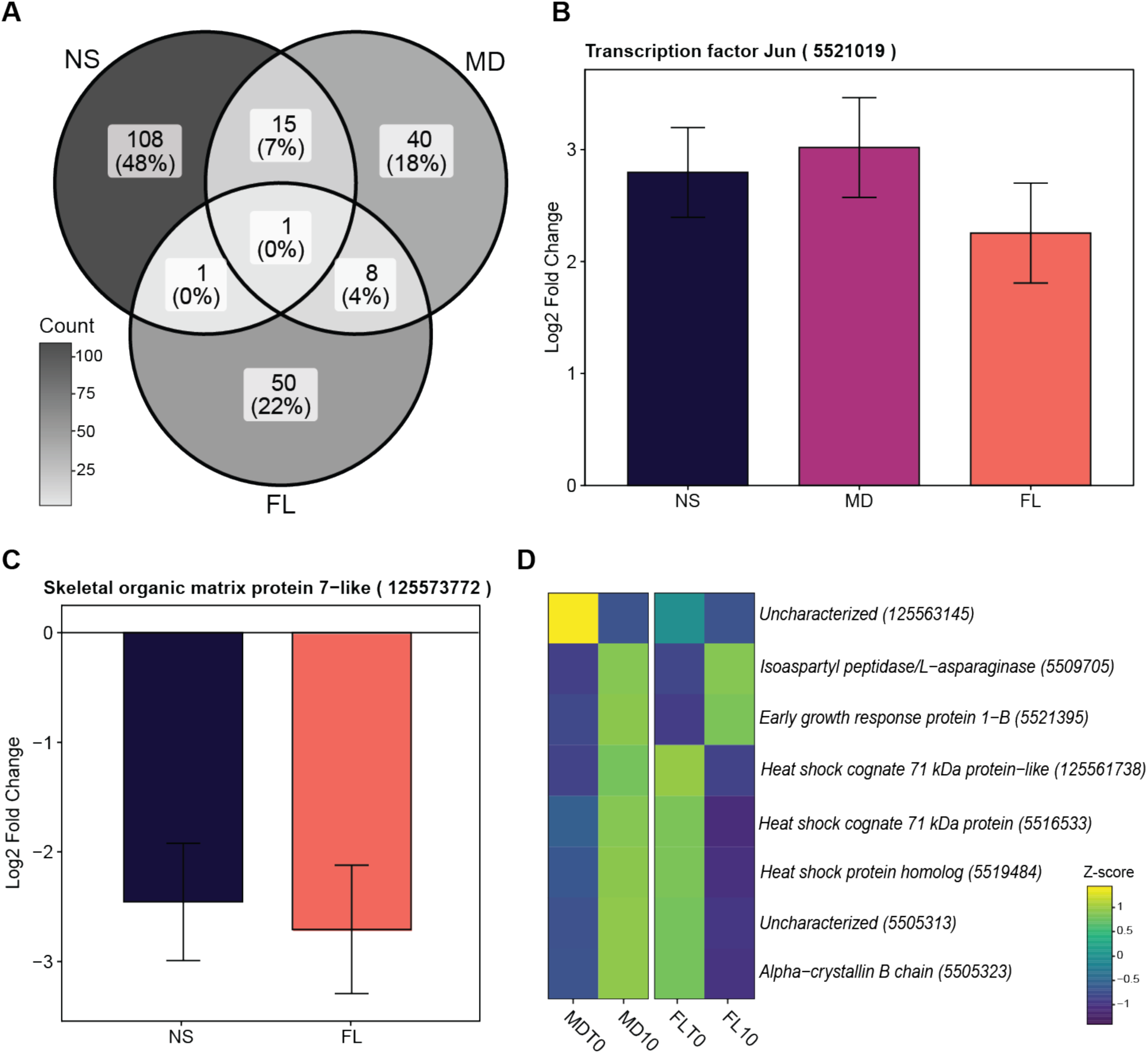
Response to cold stress in *Nematostella vectensis*. (**A**) Venn diagram depicting the shared and unique differentially expressed genes (DEGs) in cold stress (10 °C) treatments relative to timepoint zero (T0) samples. Nova Scotia (NS) on the left, Maryland (MD) on the right, and Florida (FL) on the bottom. (**B**) Gene expression of the transcription factor Jun visualized as the log2 fold change for each population. Error bars show the standard error retrieved from DESeq2 results (lfcSE). (**C**) Gene expression of the skeletal organic matrix protein 7-like gene visualized as the log2 fold change for each population. MD not shown as this gene was not a significant DEG in that population. (**D**) Heatmap visualizing z-scores of shared DEGs between MD and FL under cold stress.

### 3.4 Changes in gene expression post-heat stress include a core response and population-specific signatures

Exposure to 38 °C induced a large transcriptomic shift of nearly 10% of all genes, with NS exhibiting the largest set of significantly differentially expressed genes (n = 2539), followed by FL (n = 2380) and MD (n = 1613) (Figs. 2D-F and 4A). Comparison of these DEGs revealed a core set of 472 genes that were significantly altered in all three groups. A subset of 318 genes were co-upregulated in FL, MD, and NS (Figs. 4B and 5A), representing a conserved core thermal stress response. This core included well-known molecular defense genes such as NF-κB, NFE2L2, DnaJ, and Bcl-2 (Fig. 5B-E). Functional enrichment analysis of these co-upregulated genes confirmed a significant enrichment for protein folding (Table S2). We also identified a set of 153 genes that were significantly downregulated under heat stress across all three populations (Figs. 4C). No significant functional enrichment was detected within this gene set.

**Figure 4.**
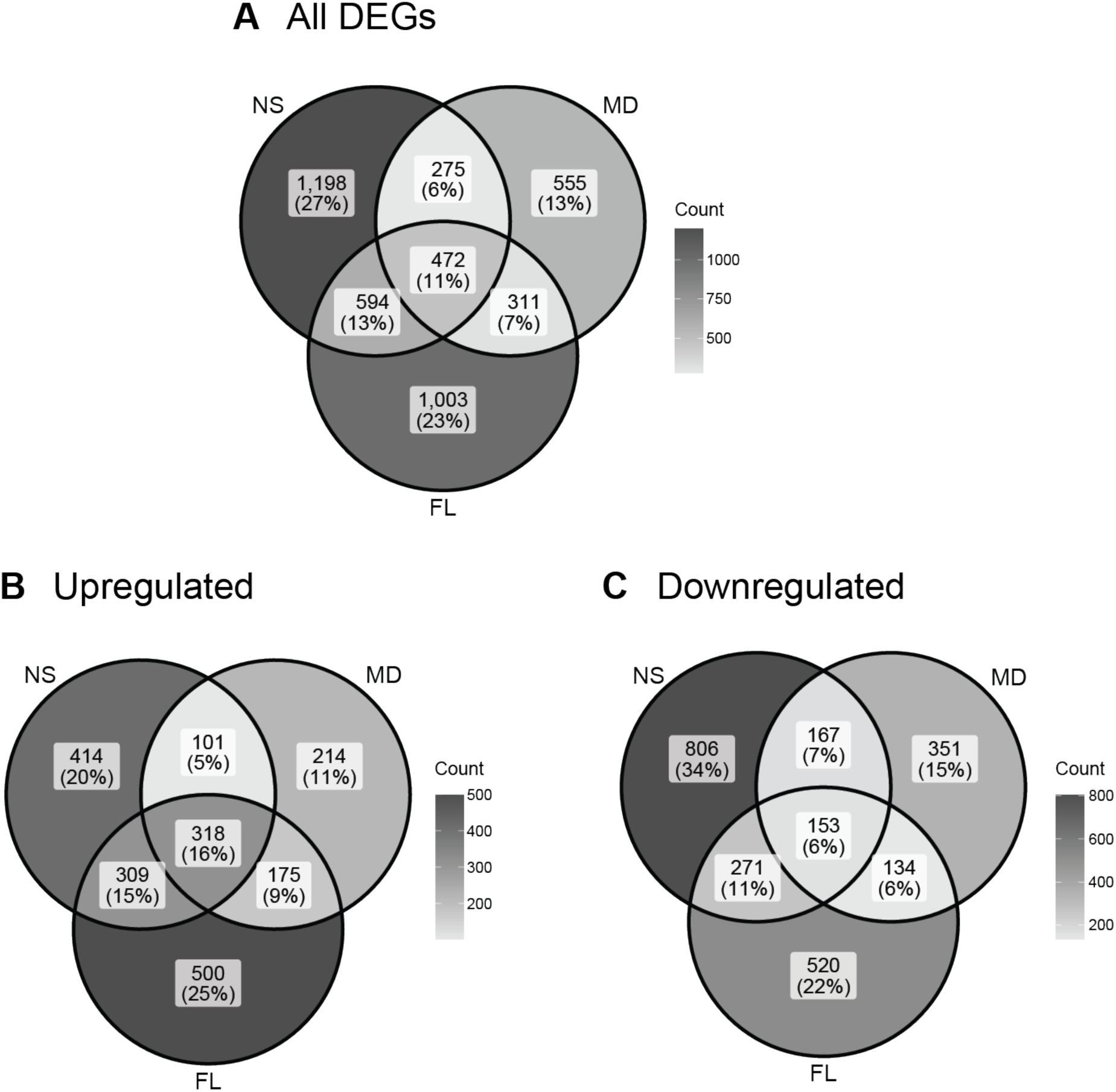
Venn diagrams of differentially expressed genes (DEGs) in heat stress (38 °C) treatments relative to timepoint zero samples (T0). (**A**) All DEGs with absolute log2 fold change > 2 and padj < 0.05. (**B**) Upregulated DEGs with log2 fold change > 2 and padj < 0.05. (**C**) Downregulated DEGs with log2 fold change < -2 and padj < 0.05.

**Figure 5.**
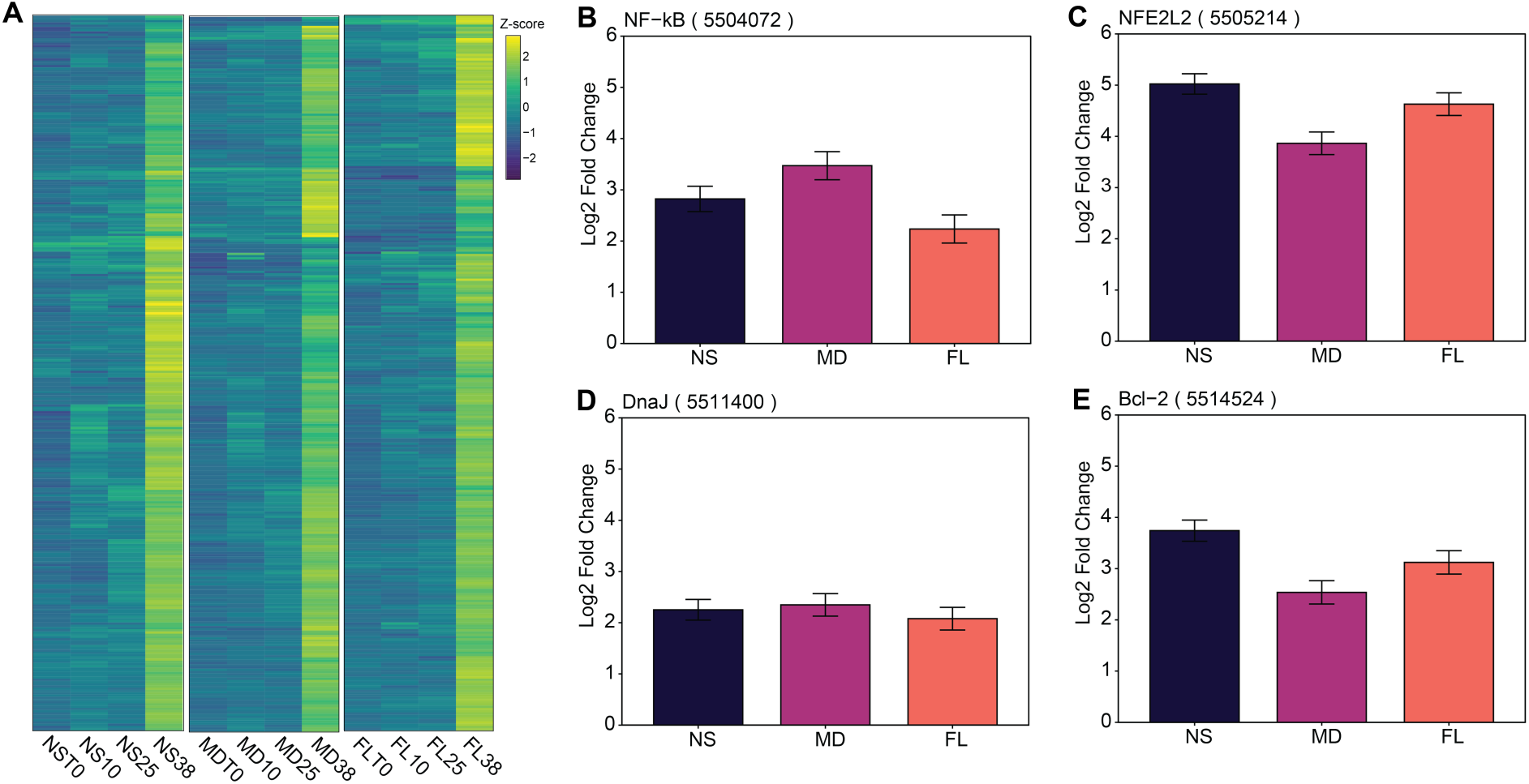
Upregulated genes under heat stress in all three *N. vectensis* populations. (**A**) Heatmap displaying z-scores of commonly upregulated genes across the three *N. vectensis* populations under heat stress. Gene expression of some example genes visualized as the log2 fold change relative to timepoint zero (T0) samples, with error bars showing the standard error retrieved from DESeq2 results (lfcSE): (**B**) NF-κB expression, (**C**) NFE2L2 expression, (**D**) DnaJ expression, and (**E**) Bcl-2 expression.

In contrast to the shared stress core, we identified a substantial number of population-specific DEGs, suggesting divergent local responses to heat. NS exhibited the most distinct response, with 1,198 genes uniquely altered. Among these, 414 uniquely upregulated genes were enriched for proteolysis, catabolic and metabolic processes, and digestion (Table S3). The 806 genes uniquely downregulated in NS were primarily associated with DNA integration and signal transduction (Table S4).

The FL population exhibited 1,003 unique DEGs, consisting of 500 uniquely upregulated and 520 uniquely downregulated genes. While the full set of 500 upregulated genes yielded no significant functional enrichment, we performed a more targeted analysis by extracting the top 100 DEGs from each population based on their log2 fold change values. From this top-expressed gene dataset, we identified a subset of 50 genes uniquely upregulated in FL (Fig. S3). Functional enrichment analysis of these top DEGs revealed a significant enrichment for nucleosome assembly (Table S5). Similarly, we identified a subset of 80 top uniquely downregulated genes in FL, which contained many DNA integration genes (Table S6).

The MD population had the fewest number of unique DEGs in response to high temperature, with only 555 population-specific genes differentially expressed. Although the full sets of 214 upregulated and 351 downregulated genes showed no significant functional enrichment, the top uniquely upregulated genes were significantly enriched for protein processing in the endoplasmic reticulum pathway (Table S7). This pathway featured key stress-response genes, including ATF-4 (116611673), dnaJ homolog subfamily C member 3 (5503853), and hypoxia up-regulated protein 1 (5506606). Additionally, the top uniquely downregulated MD genes were enriched for transcription initiation and mismatch repair processes (Table S8). Collectively, these findings suggest that while a conserved stress machinery exists in *N. vectensis*, it is overlaid by highly distinct, population-specific molecular strategies to combat heat stress.

### 3.5 DNA integration genes contribute to overall gene expression patterns under heat stress

Notably, both the NS downregulated genes as well as the top FL downregulated genes had multiple genes belonging to the DNA integration (GO:0015074) functional category. This shared functional signature was particularly intriguing given that it emerged from gene sets otherwise unique to each population. To further investigate this pattern, we analyzed the expression of all 330 genes in the *N. vectensis* genome annotated for DNA integration that exhibited a variance greater than zero across all samples. Broadly, these genes displayed a mosaic of shared and population-specific expression patterns (Fig. 6A). After filtering for uncharacterized genes, a set of 50 genes—predominantly composed of kinases, transposons, methyltransferases, and various zinc-finger domain proteins—was identified as some of the genes driving these signatures (Fig. 6B).

**Figure 6.**
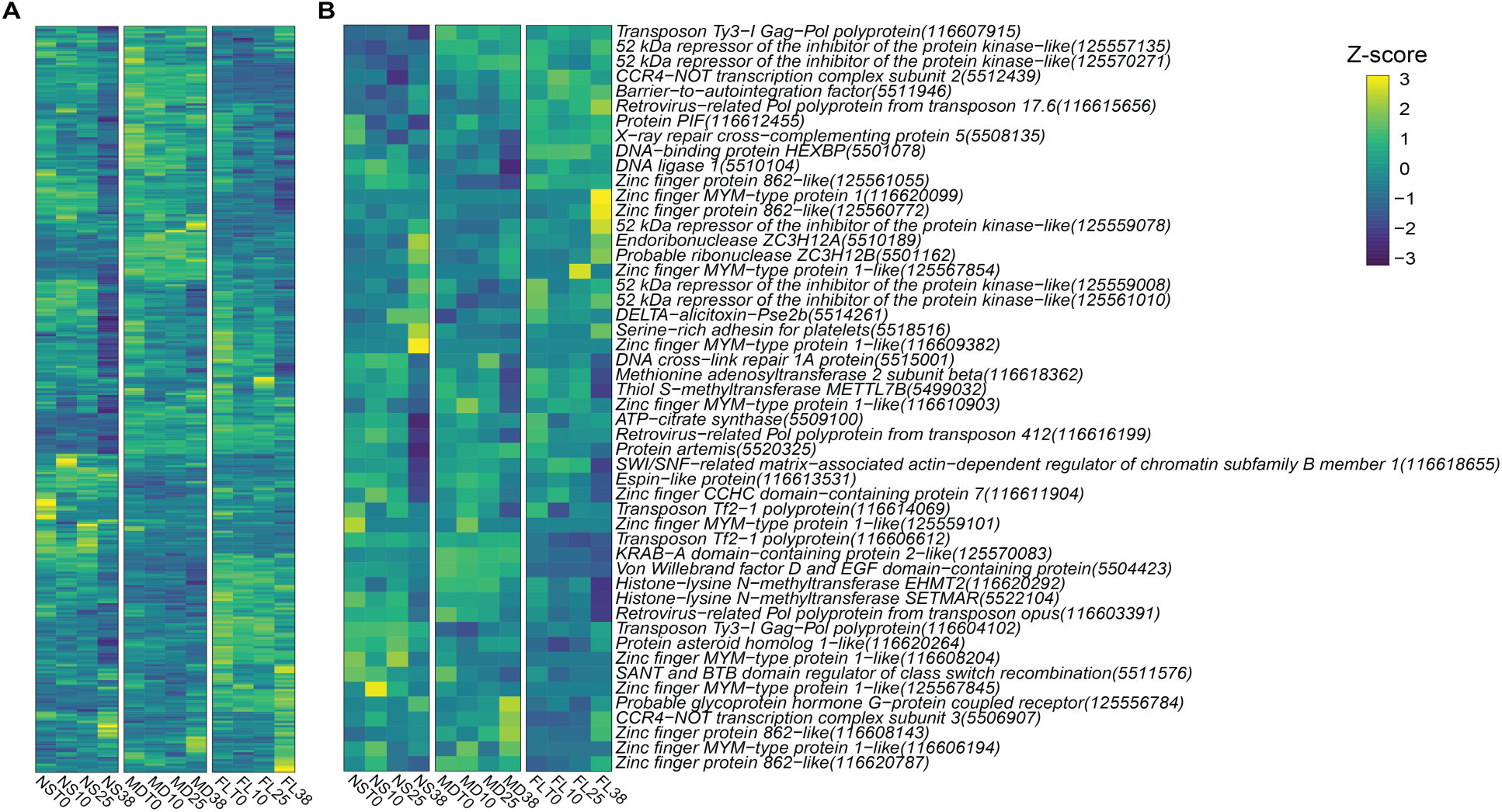
Expression patterns of DNA integration genes under heat stress. (**A**) Heatmap displaying z-scores of all 330 genes annotated with the gene ontology (GO) term ‘DNA integration’ (GO:0015074) in *N. vectensis*. (**B**) Heatmap displaying z-scores of the 50 characterized genes belonging to the DNA integration GO category.

### 3.6 Expression of the heat shock response pathway under heat stress

Given the central role of the heat shock response (HSR) in thermal tolerance, we specifically examined the expression patterns of major molecular chaperones and their primary regulator, heat shock factor 1 (HSF1), across all three populations. While all populations upregulated key HSR components by similar magnitudes (log2 fold changes depicted in Fig. 7), the MD population exhibited significantly lower normalized read counts for a key temperature-responses HSPs, including HSP70A1A, HSP90A1, and HSF1 (Bhalodi, et al. 2026; Bhalodi and Reitzel, unpublished data) compared to both FL and NS at 38 °C (Fig. S4).

**Figure 7.**
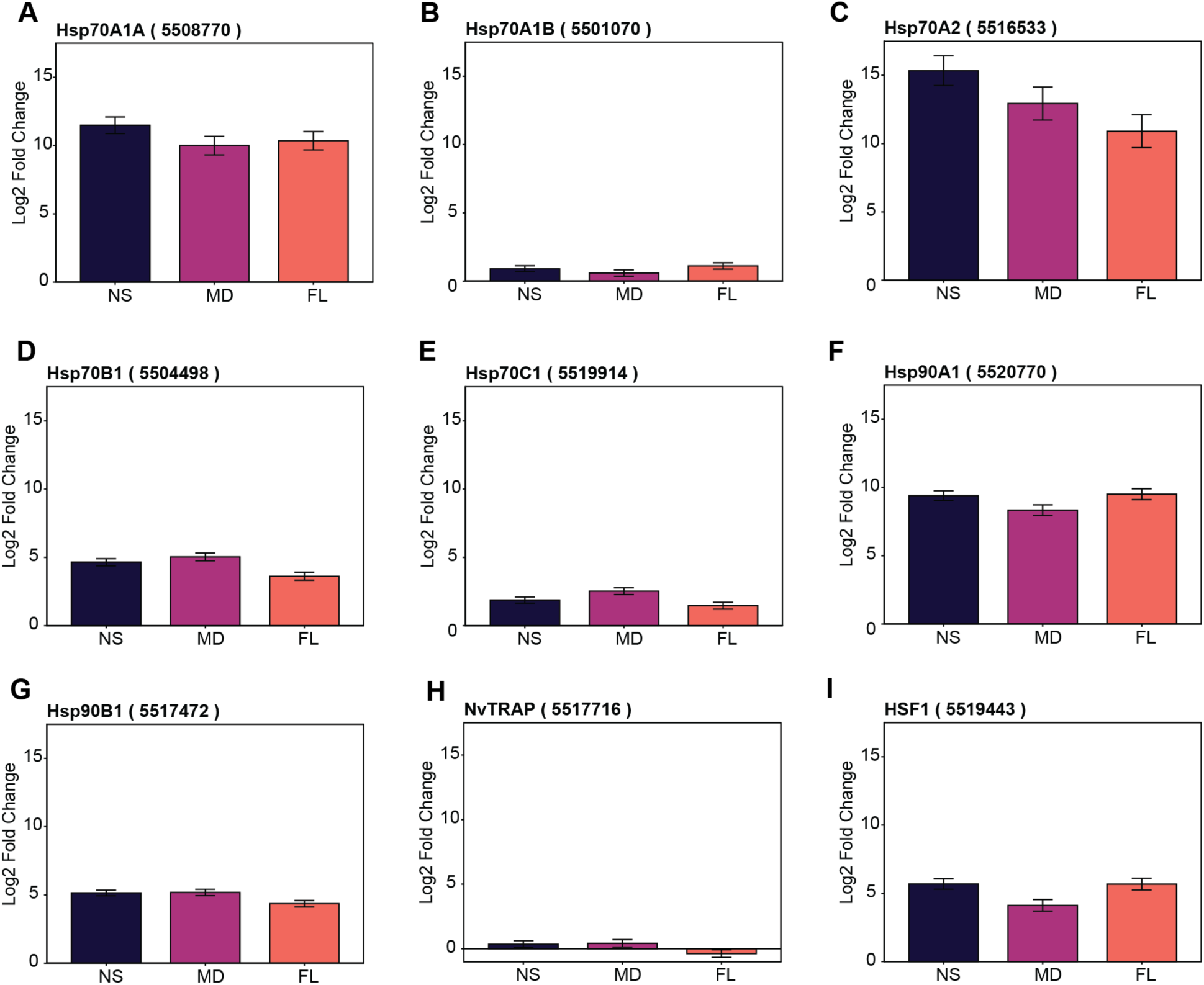
Heat shock response pathway gene expression under heat stress across *N. vectensis* populations. Gene expression visualized as the log2 fold change relative to timepoint zero (T0) samples. Error bars show the standard error retrieved from DESeq2 results (lfcSE). (**A**) Hsp70A1A expression. (**B**) Hsp70A1B expression. (**C**) Hsp70A2 expression. (**D**) Hsp70B1 expression. (**E**) Hsp70C1 expression. (**F**) Hsp90A1 expression. (**G**) Hsp90B1 expression. (**H**) NvTRAP expression. (**I**) HSF1 expression.

The presence of HSF binding sites called heat shock elements (HSEs) in gene promoters, especially a higher number of HSEs, has repeatedly been shown to be important for the regulation of genes during heat stress (Xiao, et al. 1991; Cunniff and Morgan 1993; Erkine, et al. 1999; Mendillo, et al. 2012; Ortner, et al. 2015; Li, et al. 2016; Schmauder, et al. 2022). We leveraged this characteristic to explore if highly induced heat stress genes could be targets of HSF, or if they implicate a major involvement of alternative stress pathways in *N. vectensis*. To achieve this goal, we investigated the correlation between differential gene expression and HSE counts within the putative promoters of the top 100 DEGs in each population. In both MD and FL, DEGs containing three or more HSEs were, on average, more highly upregulated than those with fewer than three (Figs. 8A–C). In contrast, the top 100 upregulated genes in NS contained no transcripts with more than four HSEs, and those with three HSEs did not show the robust induction observed in the other populations. Conversely, the majority of top-downregulated DEGs across all populations possessed fewer than two HSEs and showed no clear trend between HSE count and expression magnitude (Figs. 8D–F). Many genes with zero HSEs were strongly downregulated, suggesting that while higher HSE density drives upregulation, strong downregulation is likely governed by alternative transcription factors. These results suggest that while the MD population shows lower normalized counts for canonical HSF pathway genes, its top upregulated gene set includes multiple putative HSF targets with strong induction, supporting a hypothesis for a functional role for HSF regulation.

**Figure 8.**
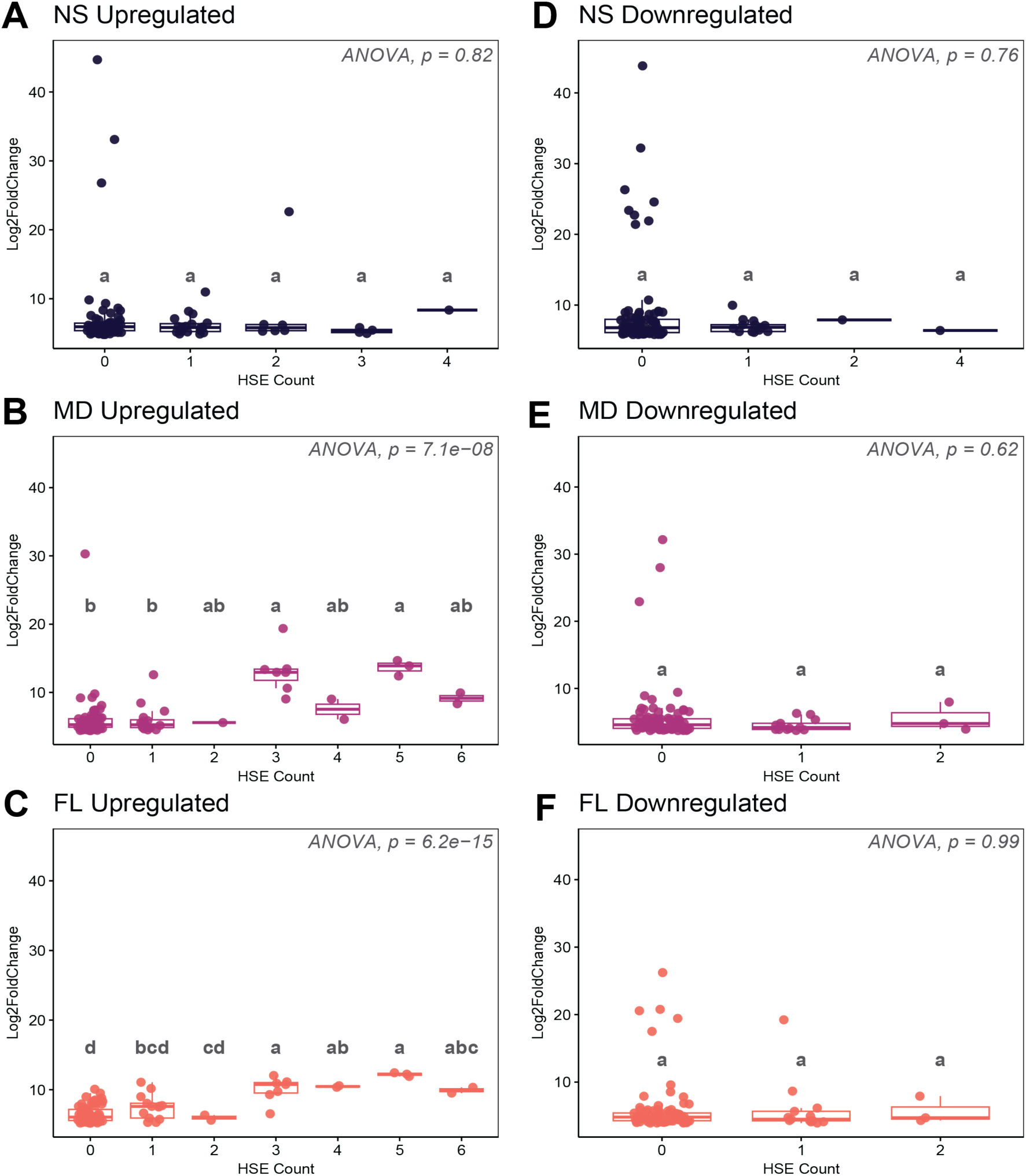
Relationship between heat shock element (HSE) abundance and magnitude of expression at 38 °C. One-way ANOVA conducted on mean log2 fold change values from each HSE category. Letters on top of boxes indicate statistical significance calculated with a post-hoc Tukey HSD test, where groups harboring unique letters represent statistically significant differences between means (p < 0.05). (**A**) Top 100 upregulated genes in Nova Scotia (NS). (**B**) Top 100 upregulated genes in Maryland (MD). (**C**) Top 100 upregulated genes in Florida (FL). (**D**) Top 100 downregulated genes in NS. (**E**) Top 100 downregulated genes in MD. (**F**) Top 100 downregulated genes in FL.

In NS, however, the top upregulated genes lacked highly induced, high-HSE targets. We hypothesized that some high-HSE genes might have been excluded from this heat stress analysis due to their presence in the 25 °C DEG dataset (see Results section 2). Indeed, we identified three major inducible chaperones—Hsp70A1A, Hsp70A2, and Hsp90A1—that overlapped with the 25 °C dataset. Intriguingly, these represent three of the only four HSPs in *N. vectensis* known to possess HSEs in their promoter regions (Bhalodi, et al. 2026). This suggests that 25 °C may constitute a physiologically stressful temperature for the NS population, which inhabits cooler environments than its southern counterparts (Sachkova, et al. 2020).

### 3.7 Contribution of alternative stress pathways to heat tolerance

Involvement of alternative stress pathways to thermal tolerance was investigated using the expression patterns of 60 genes annotated with the GO term ‘response to stress’ (GO:0006950). Many of these exhibited highly conserved expression patterns across all three populations, including heat shock proteins, MAP kinases, and ER protein processing genes such as derlin and BiP (Fig. 9). However, several genes displayed unique expression profiles that underscore the population-specific nature of the heat response. Notably, our analyses showed differences in expression of a subset of genes hypothesized to be involved in circadian biology. Cryptochrome DASH (5502007) was significantly downregulated in the FL population while remaining nearly unchanged in NS and MD. Divergent patterns were also observed among other cryptochrome-related genes. A specific cryptochrome-1 (5516268) was upregulated in both MD and FL but exhibited consistently low expression in NS at both T0 and post-heat treatment. In contrast, cryptochrome-2 (5510586) was upregulated in NS and MD but showed minimal induction in FL. Interestingly, not all members of this family diverged; a separate cryptochrome-1 gene (5504485) was consistently downregulated across all three populations. These results suggest that while many core cellular stress genes are similarly utilized by each population, the regulation of some genes, particularly the cryptochromes, have been fine-tuned differently across these populations.

**Figure 9.**
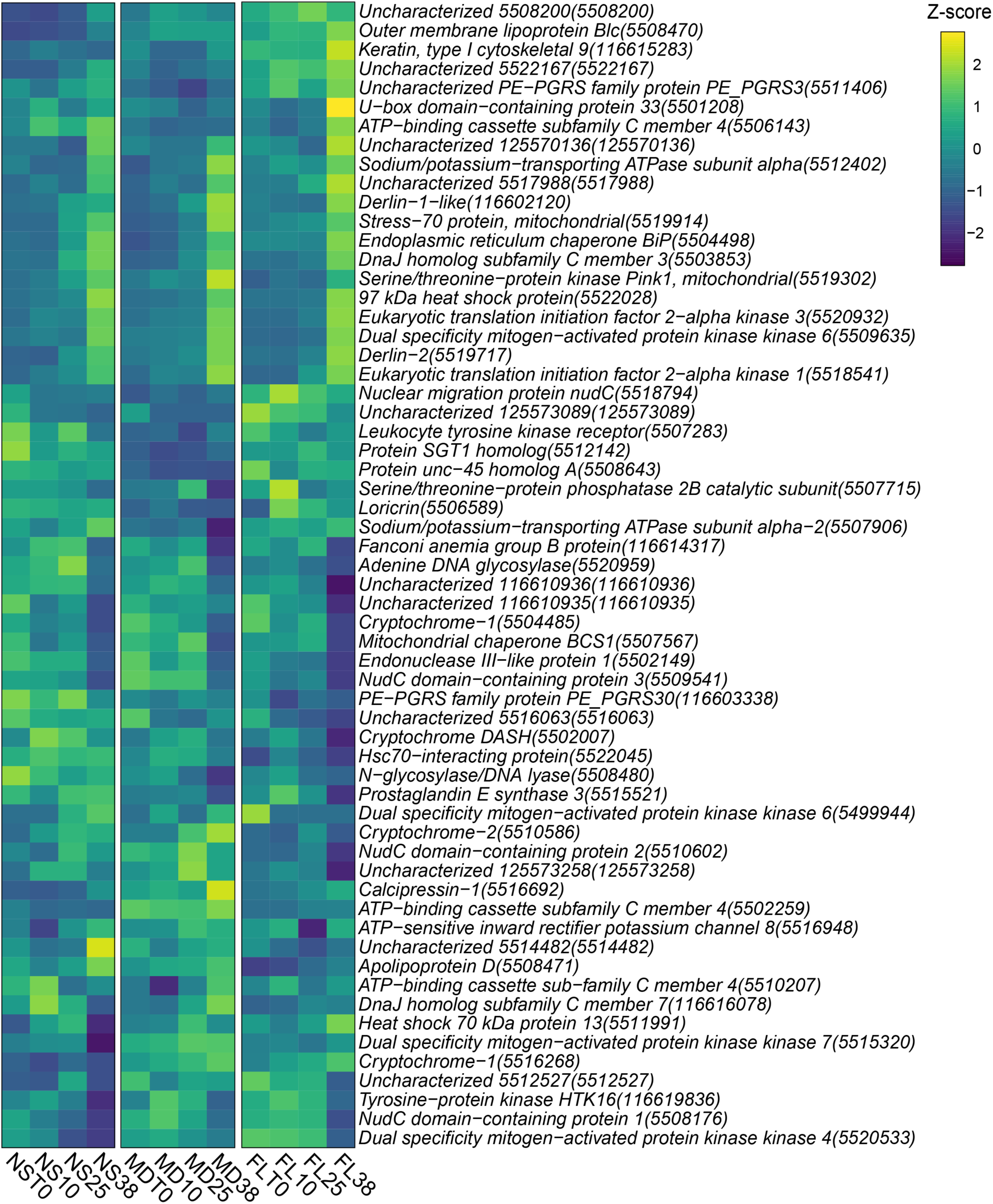
Heatmap displaying z-scores of 60 genes annotated with the gene ontology (GO) term ‘response to stress’ (GO:0006950) in *N. vectensis*.

### 3.8 Diverse transcriptional control of heat stress responses across populations

The presence of population-specific gene expression patterns to heat stress prompted us to explore whether unique transcription factors might be regulating these responses across individual populations. To identify the transcription factors driving the patterns of gene expression, we searched for enriched binding motifs within the putative promoters of the top 100 up- and down-regulated genes in each population. In several instances, the analysis yielded broad or ambiguous results. For example, a motif characterized by alternating cytosine and adenine residues was enriched in the NS upregulated set; while this motif matches various zinc finger domains in database searches, the functional diversity and abundance of zinc finger proteins make specific biological interpretation challenging (Fig. S5). Similar results were obtained for the MD downregulated set, and no significant motifs were identified for the FL downregulated genes.

However, more specific regulatory signatures emerged from some of the comparisons. Interestingly, HSEs were significantly enriched only within the FL upregulated gene set (Fig. 10). Furthermore, we identified an enrichment for the cAMP-response element-binding protein (CREB) transcription factor binding site in the upregulated gene sets of both FL and MD. In contrast, the homeobox domain binding site was the top match for the motif enriched within the NS downregulated genes. These findings suggest that the molecular response to heat stress in *N. vectensis* may be governed by distinct transcription factors among populations, providing evidence for potential local adaptation in shaping the regulatory landscape of thermal tolerance.

**Figure 10.**
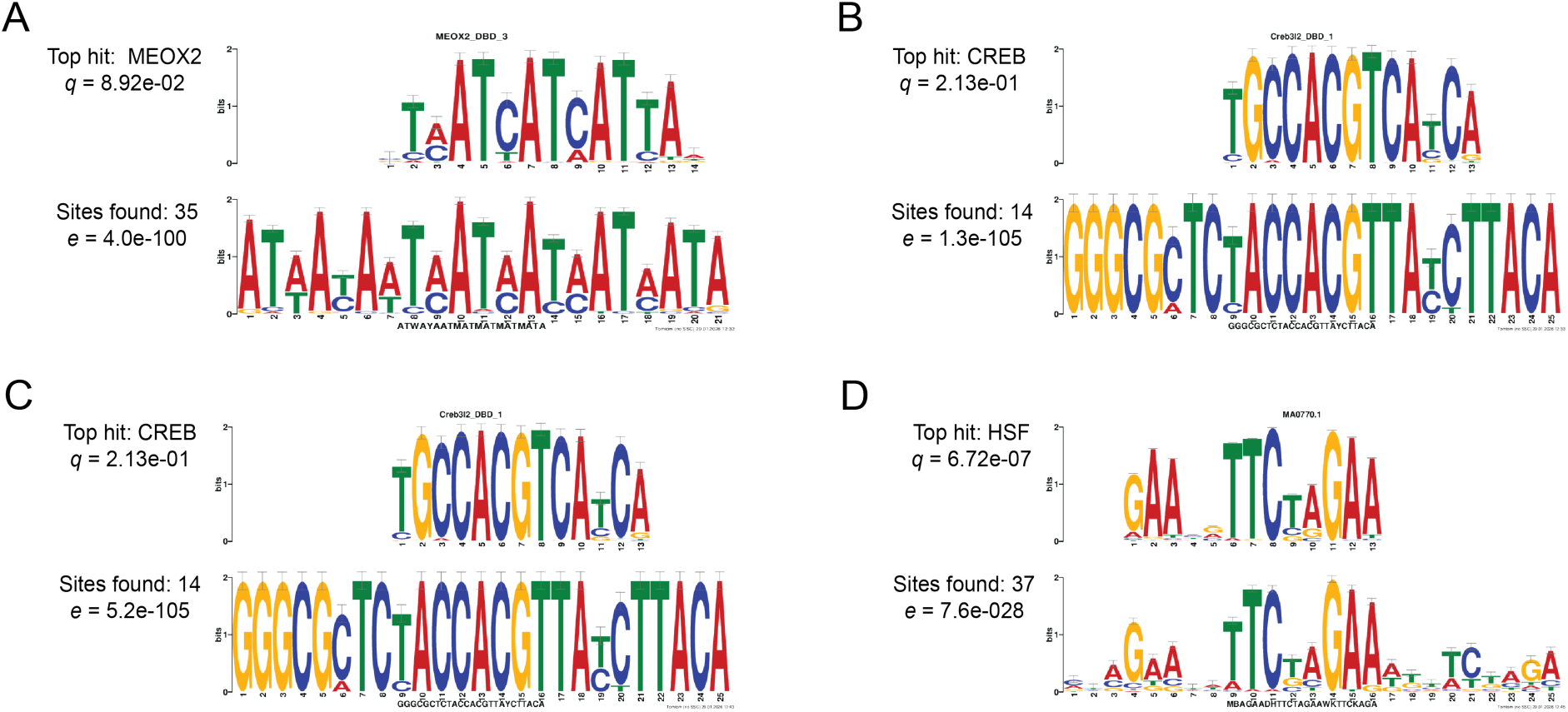
Motif enrichment analysis in differentially expressed genes under heat stress. In each case, the bottom motif is an enriched motif identified in a set of putative promoter sequences using the MEME program from MEME Suite (https://meme-suite.org/meme/). The top motif is the top hit from popular transcription factor binding motif databases that match the enriched motif, identified using the TOMTOM program from MEME Suite. Motif enrichment for: (**A**) Top 100 downregulated genes in Nova Scotia (NS), (**B**) Top 100 upregulated genes in Maryland (MD), (**C**) Top 100 upregulated genes in Florida (FL), and (**D**) Top 100 upregulated genes in FL.

## 4. Discussion

We used RNA-seq to examine gene expression in populations of the sea anemone *Nematostella vectensis* exposed to acute cold (10 °C) and heat (38 °C) stress. By using clonal populations from Nova Scotia (NS), Maryland (MD), and Florida (FL), we mitigated the impact of individual genetic variation within populations while exploring the transcriptional response diversity of genetically distinct populations. Broadly, our results revealed limited gene expression changes in response to cold and a robust response to heat (Fig. 2). This asymmetrical transcription response to an identical change in absolute temperature (i.e., ± 13°C) differs from earlier studies in the temperate coral *Astrangia poculata*, which exhibited a significantly larger response to cold (4–6 °C) than to heat (30–31 °C) (Wuitchik, et al. 2021; Wuitchik, et al. 2024). However, those corals were exposed to comparatively lower temperatures for a longer period (15–16 days), suggesting that extended exposure to lower experimental temperatures may elicit a stronger cold response.

The natural habitat range of *N. vectensis* allowed us to investigate whether observed gene expression patterns corresponded to the climates these populations naturally experience; NS and FL reside in relatively cold versus warm habitats, respectively, while MD experiences an intermediate climate. Our heat stress data reveal a larger transcriptional response in the range-extreme populations (NS and FL) compared to the intermediate MD population (Figs. 2 and 4A). This finding aligns with the Climate Variability Hypothesis (CVH), which posits that organisms experiencing high seasonal variability possess higher plasticity, contributing to a broader range of thermal tolerances. Populations inhabiting the extremes of a species’ range are likely to experience highly variable climates, which suggests that NS and FL may have evolved more plastic and, therefore, tolerant responses to a wider range of temperatures. While the CVH would predict similar patterns in the cold, MD exhibited a response to cold comparable to that of the FL population. This exception may be explained by the longer duration that the MD population has been in laboratory culture (40-50 years), potentially making it more adapted to the narrower temperature ranges in the lab environment than the other field-collected populations.

Gene Ontology (GO) analyses revealed that genes involved in maintaining proteostasis were among the most commonly differentially expressed across both temperature treatments and all populations. These included genes belonging to the heat shock protein (HSP) families, their regulators such as HSF1, and co-chaperones such as DnaJ. HSPs are highly conserved molecular chaperones that maintain the integrity of proteins by facilitating protein folding and preventing aggregation in the cell (Lindquist 1986). DnaJ proteins stabilize the interaction of HSPs and their substrate proteins, thereby acting as crucial partners of Hsp70 (Qiu, et al. 2006). The transcription of HSPs is governed by the HSF family of proteins, which are also responsible for activating other genes that help mitigate the impact of cellular stress (Santoro 2000). The expression patterns of these genes were population-specific under cold stress; Many of the shared cold stress DEGs between FL and MD belonged to this category, yet they displayed opposing expression patterns between the two populations (Fig. 3D). Specifically, FL upregulated these genes while MD downregulated them, indicating that FL anemones may have a greater need for maintaining protein integrity at this temperature. Interestingly, many of these proteostasis-related genes showed elevated expression in the NS population at the control temperature (25 °C), suggesting that this temperature may already induce some proteotoxicity for these higher latitude locations. Previous research has shown an optimal average growth rate for this population at 21 °C (Reitzel, Chu, et al. 2013) and higher mortality in NS anemones cultured at 25 °C (Baldassarre, et al. 2023), further supporting the hypothesis that our control temperature may have induced some physiological stress for the NS anemones.

Our analyses also revealed a distinct signature of heat stress expression for genes belonging to the “DNA integration” Gene Ontology category (GO:0015074), which encompasses genes that facilitate the incorporation of a DNA segment into another, often larger DNA sequence. We observed significant enrichment for this GO term in the uniquely downregulated gene sets of both the NS and FL populations. This indicates that while both populations downregulate DNA integration processes, they are, at least in part, recruiting distinct gene sets to achieve this outcome. RNA-seq studies conducted on stress responses in other cnidarians have similarly shown an overrepresentation of genes in the DNA integration category (Hou, et al. 2018; Vidal-Dupiol, et al. 2022). This prompted us to explore the expression of 330 identified DNA integration genes more broadly, revealing a mosaic of shared and unique patterns; while some responses are conserved across populations, others exhibit expression profiles unique to specific groups (Fig. 6A). While this gene set was largely dominated by uncharacterized transcripts, the 50 annotated genes—comprising methyltransferases, transposable elements (TEs), and zinc-finger proteins—did not reveal any distinct functional patterns across the populations (Fig. 6B). Nonetheless, transposable elements play a critical role in genome organization and function (Feschotte 2008), contributing to rearrangements or expansions that influence regulatory architecture and gene expression (Casacuberta and González 2013). Transposon-related genes are known to be upregulated under high seawater CO2 concentrations in the sea anemone *Anemonia viridis* (Urbarova, et al. 2019), under heat stress in the coral *Acropora hyacinthus* (Rose, et al. 2015), and during white plague disease infection in the holobiont of the coral *Orbicella faveolata* (Daniels, et al. 2015). Intriguingly, research has identified the promoters of *Drosophila* HSP genes as targets for evolution by transposable elements (Walser, et al. 2006), thereby driving diversity in the regulation of such genes. Additionally, previous research in *Caenorhabditis* species has demonstrated a disproportionately high density of HSEs within helitron transposable elements; these HSEs contribute to the diversification of the heat shock response in *C. elegans* via transposon-mediated mobilization (Garrigues, et al. 2019). While mobile elements may offer increased genetic diversity and potentially enhance stress tolerance via the acquisition of regulatory elements and increased expression of proteostasis genes, their activity may also pose a threat to genomic stability. Notably, the DNA methylation machinery in cnidarians has been shown to specifically target transposons, thereby protecting the genome against the harmful activity of mobilizing elements (Ying, et al. 2022). Although our data did not suggest an obvious pattern of upregulation in the methylation machinery itself, the mechanistic link between mobile elements, methylation, and cellular stress response presents a promising avenue for future research.

Examination of genes belonging to the “response to stress” GO category (GO:0006950) also revealed shared and unique expression patterns under heat stress across populations (Fig. 9). Of particular interest was the apparent “fine-tuning” observed in the expression of various cryptochrome genes. For instance, we observed significant downregulation of cryptochrome DASH (5502007) in the FL population, whereas its expression remained largely unchanged in the others. Because cryptochromes are light-responsive, circadian clock-related genes (Leach and Reitzel 2020), their differential expression was unexpected given that our experiment was conducted entirely in the dark. Nonetheless, cryptochromes are also known to modulate innate immunity and inflammatory responses, acting as molecular links between the circadian clock and stress pathways (Baxter and Ray 2020). Accordingly, changes in cryptochrome expression have been documented in response to heat stress in the *Acropora tenuis-Cladocopium* sp. holobiont (Gong, et al. 2024), and during thermal and nutrient stress in the coral *Acropora aspera* (Rosic, et al. 2014). However, the specific mechanisms by which these genes contribute to population-specific transcriptional responses at the molecular level require further investigation.

Lastly, our data indicate that population-specific signatures in response to heat stress may be orchestrated by distinct transcription factors in each population. Specifically, the promoters of upregulated genes in the FL population showed enrichment for cAMP response element-binding protein (CREB) and HSF transcriptional regulators. In the MD population, the promoters of upregulated genes displayed an overrepresentation of only CREB motifs, whereas homeobox motifs were enriched in the promoters of genes downregulated in NS. Like HSFs, CREBs are multifaceted transcription factors that regulate genes involved in a variety of biological processes; however, their specific role in immune function and stress responses is also well-established (Wen, et al. 2010). Homeobox domain-containing proteins are also important regulators of the stress response; for example, the FoxO family of transcription factors plays a critical role in oxidative stress resistance (Bridge, et al. 2010). Together, our findings imply that populations of *N. vectensis* exhibit unique transcriptional responses to identical stressors, which are likely facilitated by distinct transcriptional regulators.

## Acknowledgements

The authors would like to thank Dr. Auston Rutlekowski (Reitzel lab) for help with experimental setup and sample collection for this research project. Additionally, the authors would like to thank all members of the Reitzel lab for constructive comments during the development of this project and the preparation of this manuscript.

## Competing Interests

The authors declare no competing interests.

## Funding

This research was supported by NSF Award 1545539 and NIH Award R15-GM137253.

## Data and Resource Availability

All RNA sequencing data and metadata are deposited on the NCBI’s Sequence Read Archive (SRA) database under accession PRJNA1190397. Additionally, all scripts used to analyze sequence data are available on GitHub at https://github.com/jbhalodi/Temperature-Stress-RNA-Seq.

## Abbreviations

HSF: heat shock factor
T0: timepoint zero
NS: Nova Scotia
MD: Maryland
FL: Florida
PPT: parts per thousand
ASW: artificial seawater
RIN: RNA-integrity number
HSE: heat shock element
PCA: principal component analysis
DEG: differentially expressed gene
HSP: heat shock protein
ANOVA: analysis of variance
HSR: heat shock response
GO: gene ontology
CREB: cAMP-response element-binding protein
TE: transposable element
ER: endoplasmic reticulum
CVH: climate variability hypothesis

## Notes

### Competing Interest Statement

The authors have declared no competing interest.

