## Supplementary Material for "A Matter of Degrees: Latitudinal Variation in the Transcriptional Response to High and Low Temperatures in an Estuarine Cnidarian"

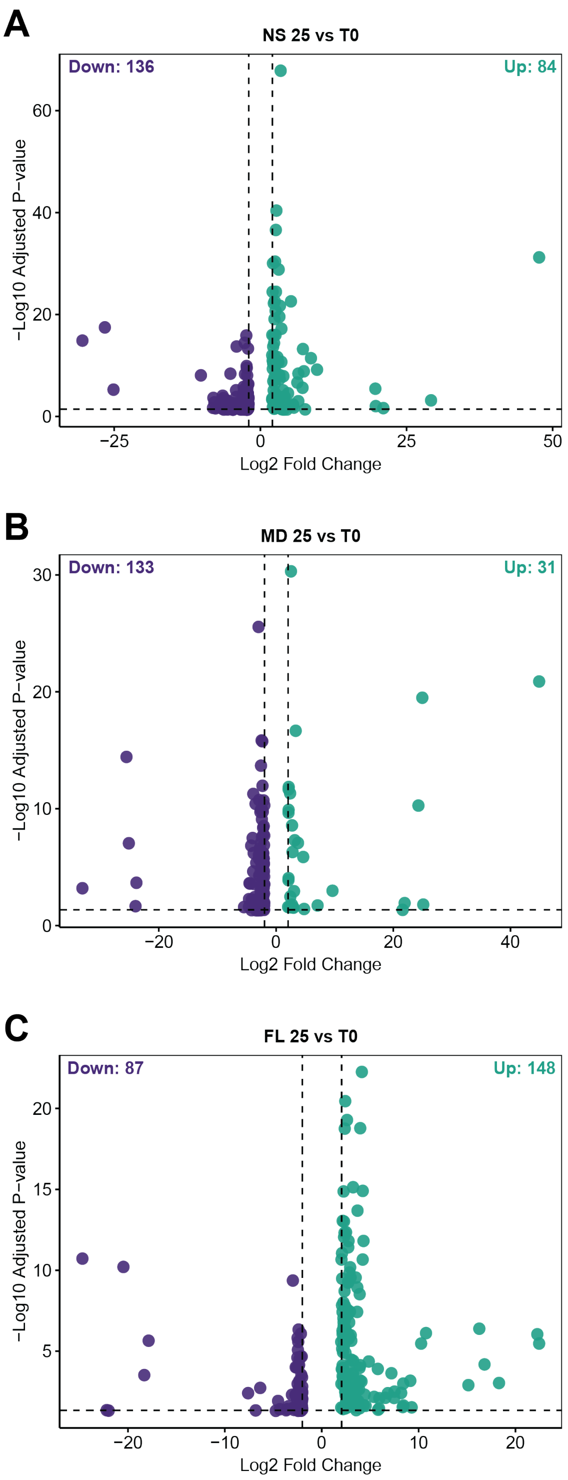


**Figure S1.** Volcano plots of differentially expressed genes in control samples (25 °C) relative to timepoint zero samples (T0). Significantly differentially downregulated genes (log2 fold change < -2, padj < 0.05) in purple (left) and upregulated genes (log2 fold change > 2, padj < 0.05) in teal (right). (**A**) Nova Scotia (NS). (**B**) Maryland (MD). (**C**) Florida (FL).

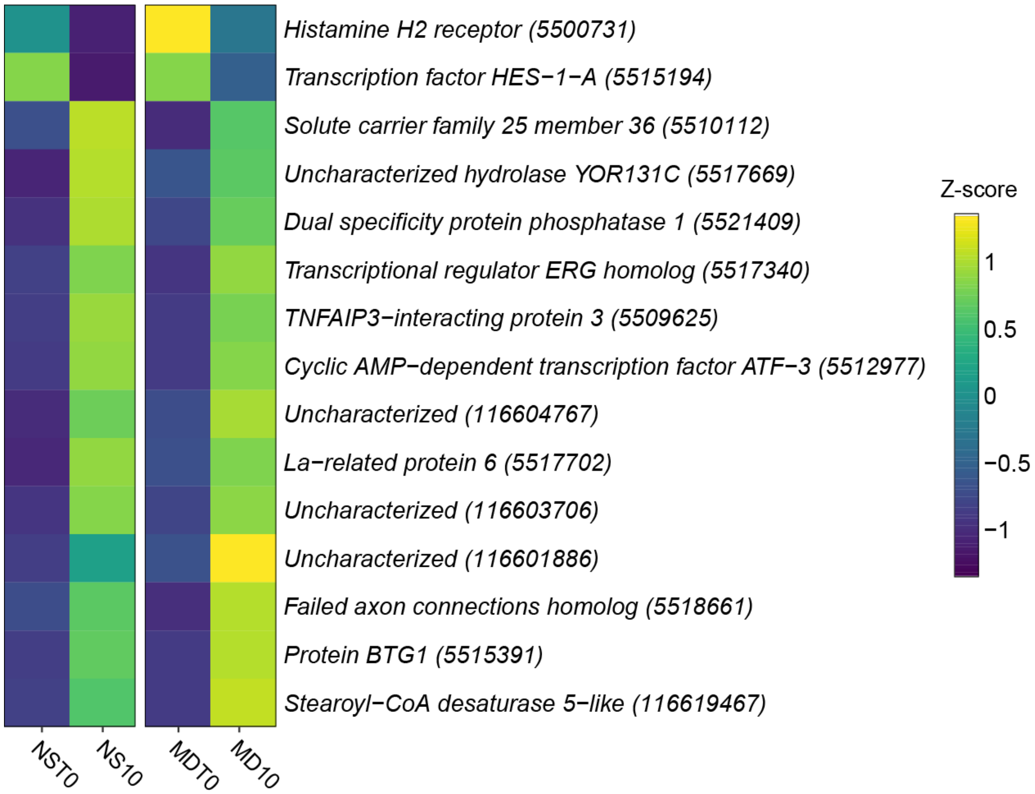


**Figure S2.** Heatmap visualizing z-scores of shared differentially expressed genes (DEGs) between Nova Scotia (NS) and Florida (FL) under cold stress (10 °C).

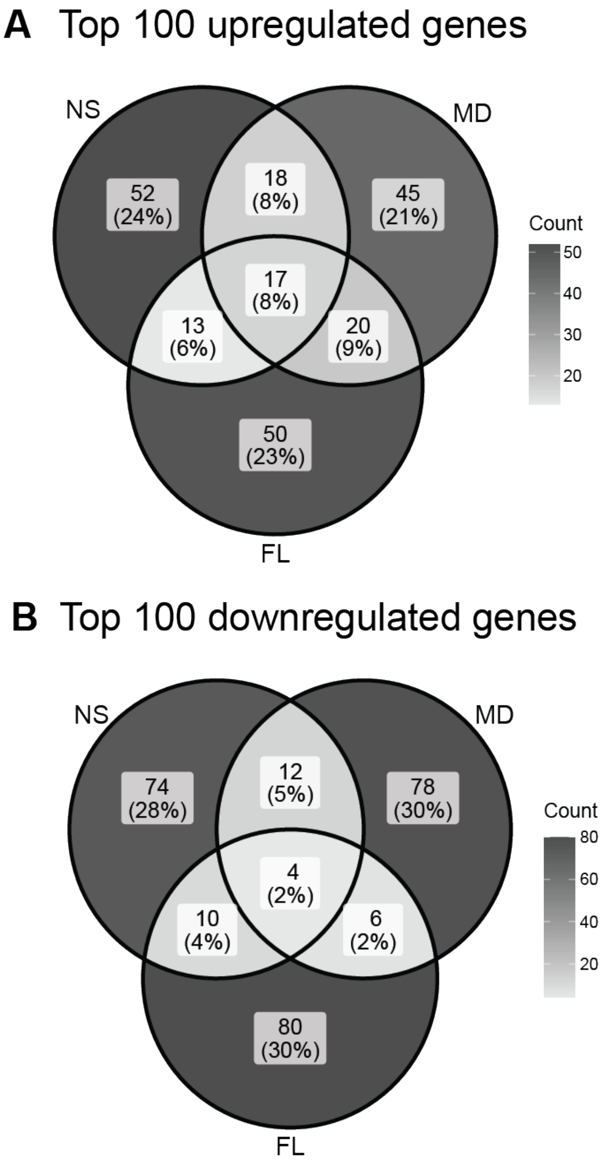


**Figure S3.** Venn diagrams of top 100 differentially expressed genes (DEGs) in heat stress (38 °C). (**A**) Top 100 upregulated DEGs with padj < 0.05. (**B**) Top 100 downregulated DEGs with padj < 0.05.

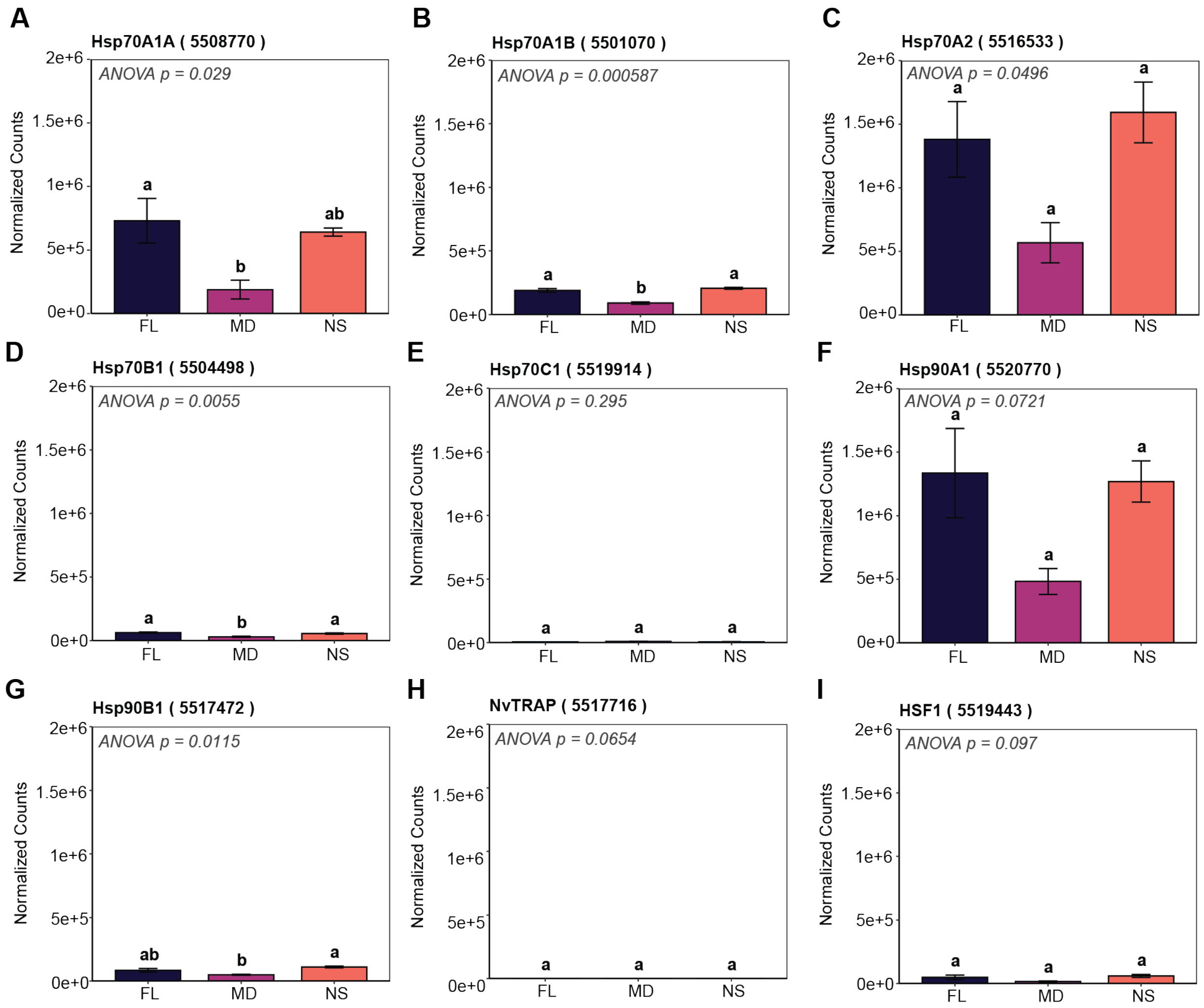


**Figure S4.** Heat shock response pathway gene expression at 38 °C across *Nematostella* populations. Gene expression visualized as the normalized read counts at 38 °C for all samples. One-way ANOVA conducted on normalized read counts. Letters on top of bars indicate statistical significance calculated with a post-hoc Tukey HSD test, where bars harboring unique letters represent statistically significant difference between means ($p < 0.05$). Error bars show the standard error of the mean. (**A**) Hsp70A1A expression. (**B**) Hsp70A1B expression. (**C**) Hsp70A2 expression. (**D**) Hsp70B1 expression. (**E**) Hsp70C1 expression. (**F**) Hsp90A1 expression. (**G**) Hsp90B1 expression. (**H**) NvTRAP expression. (**I**) HSF1 expression.

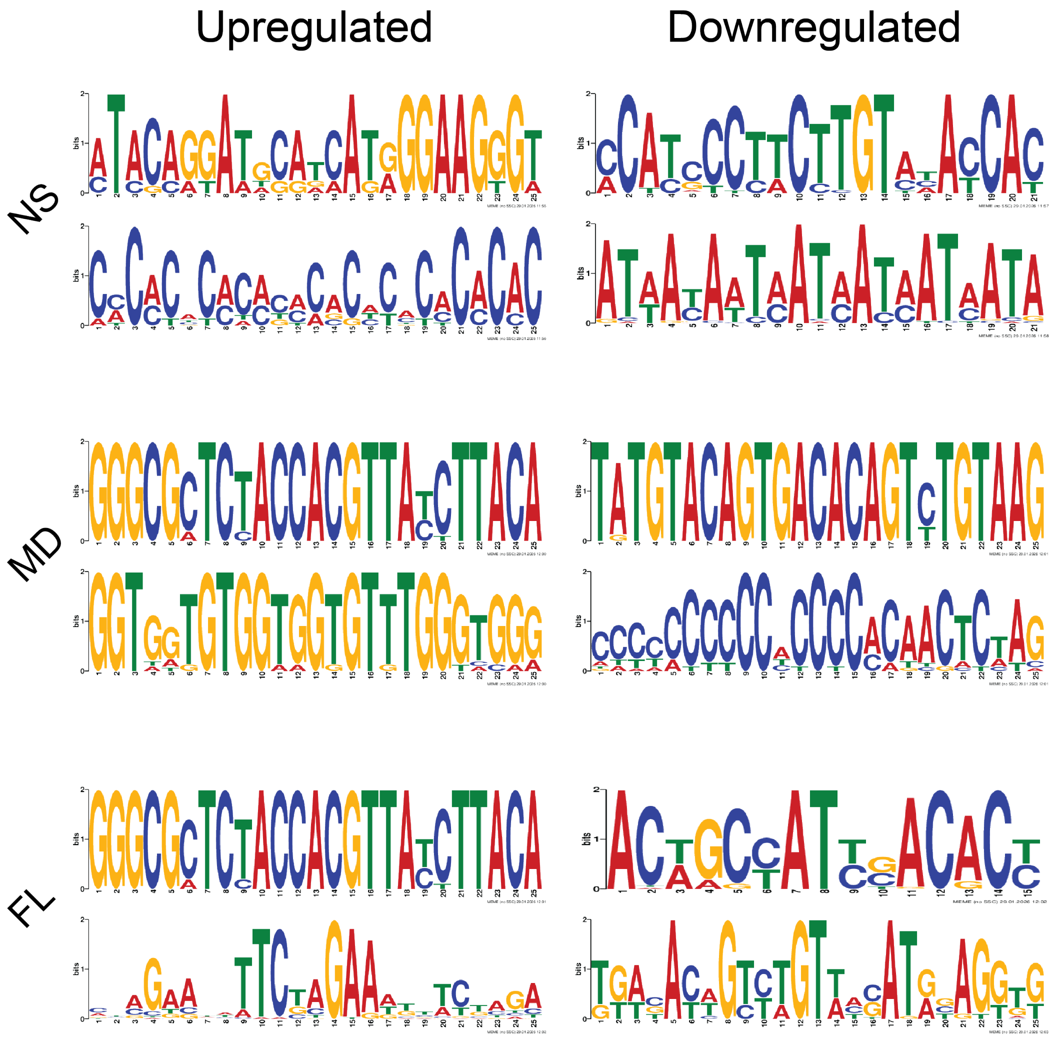


**Figure S5.** Motif enrichment analysis in differentially expressed genes under heat stress. Putative promoter sequences were used for the top 100 up- and downregulated genes from each population for enrichment analysis. Motifs are enriched motifs identified in each set of putative promoter sequences using the MEME program from MEME Suite (https://meme-suite.org/meme/).

| Term | Count | Benjamini |
|---|---|---|
| **protein refolding** | **5** | **1.45E-09** |
| **response to heat** | **3** | **8.50E-05** |
| **cellular response to unfolded protein** | **2** | **9.16E-03** |
| vesicle-mediated transport | 2 | 1.01E-01 |

**Table S1.** Gene ontology (GO) analysis of differentially expressed genes (DEGs) common in Maryland (MD) and Florida (FL) under cold stress (10 °C). Significantly enriched GO terms bolded.

| Term | Count | Benjamini |
|---|---|---|
| **protein folding** | **8** | **0.03** |
| response to endoplasmic reticulum stress | 4 | 0.11 |
| regulation of transcription by RNA polymerase II | 21 | 0.11 |
| amino acid transmembrane transport | 5 | 0.11 |
| negative regulation of MAPK cascade | 3 | 0.25 |
| innate immune response | 4 | 0.56 |
| ubiquitin-dependent protein catabolic process | 6 | 0.56 |
| protein import into mitochondrial matrix | 3 | 0.61 |
| regulation of apoptotic process | 5 | 0.71 |
| positive regulation of proteasomal ubiquitin-dependent protein catabolic process | 3 | 1.00 |
| response to unfolded protein | 2 | 1.00 |
| protein phosphorylation | 9 | 1.00 |
| modification-dependent protein catabolic process | 2 | 1.00 |

**Table S2.** Gene ontology (GO) analysis of differentially expressed genes (DEGs) common in Nova Scotia (NS), Maryland (MD), and Florida (FL) under heat stress (38 °C). Significantly enriched GO terms bolded.

| Term | Count | Benjamini |
|---|---|---|
| **proteolysis** | **44** | **1.08E-16** |
| **chitin catabolic process** | **5** | **3.67E-04** |
| **lipid catabolic process** | **8** | **2.40E-03** |
| **carbohydrate metabolic process** | **11** | **2.53E-03** |
| **digestion** | **4** | **7.69E-03** |
| **ganglioside catabolic process** | **4** | **0.02** |
| negative regulation of membrane protein ectodomain proteolysis | 3 | 0.10 |
| collagen catabolic process | 4 | 0.15 |
| arachidonate secretion | 4 | 0.18 |
| phospholipid metabolic process | 4 | 0.31 |
| extracellular matrix organization | 5 | 0.31 |
| blood coagulation | 2 | 0.48 |
| sodium ion homeostasis | 2 | 0.48 |
| chemical synaptic transmission | 8 | 0.61 |
| positive regulation of fibroblast growth factor receptor signaling pathway | 2 | 0.79 |
| chloride transmembrane transport | 4 | 0.79 |
| potassium ion homeostasis | 2 | 0.84 |
| chloride ion homeostasis | 2 | 0.84 |
| mitochondrial transport | 2 | 0.90 |
| cell volume homeostasis | 2 | 0.90 |

**Table S3.** Gene ontology (GO) analysis of upregulated genes unique to Nova Scotia (NS) under heat stress (38 °C). Significantly enriched GO terms bolded.

| Term | Count | Benjamini |
| --- | --- | --- |
| **DNA integration** | **37** | **6.59E-10** |
| **G protein-coupled receptor signaling pathway** | **66** | **6.10E-06** |
| double-strand break repair via homologous recombination | 9 | 0.14 |
| regulation of intracellular pH | 6 | 0.37 |
| DNA replication | 9 | 0.52 |
| monoatomic ion transmembrane transport | 8 | 0.70 |
| transcription preinitiation complex assembly | 6 | 1.00 |
| cellular response to light stimulus | 9 | 1.00 |
| phototransduction | 9 | 1.00 |
| DNA-templated transcription initiation | 6 | 1.00 |
| methylation | 9 | 1.00 |
| mitotic cell cycle | 6 | 1.00 |
| mitotic spindle assembly checkpoint signaling | 3 | 1.00 |
| nervous system process | 7 | 1.00 |

**Table S4.** Gene ontology (GO) analysis of downregulated genes unique to Nova Scotia (NS) under heat stress (38 °C). Significantly enriched GO terms bolded.

| Term | Count | Benjamini |
|---|---|---|
| **nucleosome assembly** | **4** | **0.01** |
| proteolysis | 4 | 0.28 |
| G protein-coupled receptor signaling pathway | 5 | 0.28 |

**Table S5.** Gene ontology (GO) analysis of top upregulated genes unique to Florida (FL) under heat stress (38 °C). Significantly enriched GO terms bolded.

| Term | Count | Benjamini |
|---|---|---|
| DNA integration | 5 | 0.07 |
| G protein-coupled receptor signaling pathway | 6 | 0.70 |

**Table S6.** Gene ontology (GO) analysis of top downregulated genes unique to Florida (FL) under heat stress (38 °C).

| Term | Count | Benjamini |
|---|---|---|
| **Protein processing in endoplasmic reticulum** | **3** | **0.05** |

**Table S7.** Gene ontology (GO) analysis of top upregulated genes unique to Maryland (MD) under heat stress (38 °C). Significantly enriched GO terms bolded.

| Term | Count | Benjamini |
|---|---|---|
| **transcription preinitiation complex assembly** | **4** | **0.01** |
| **DNA-templated transcription initiation** | **4** | **0.01** |
| **mismatch repair** | **3** | **0.03** |

**Table S8.** Gene ontology (GO) analysis of top downregulated genes unique to Maryland (MD) under heat stress (38 °C). Significantly enriched GO terms bolded.
